# Association between *KCNC1* gene polymorphism and blood lipid levels in determining the efficacy of high-intensity interval training

**DOI:** 10.1101/2025.11.23.690038

**Authors:** Junren Lai, Li Gong, Yan Liu, Yanchun Li, Jing Nie, Duoqi Zhou

## Abstract

**Objective:** The potassium voltage-gated channel subfamily C member 1 (KCNC1) gene plays a crucial role in both neural excitability and lipid metabolism. However, the impact of its genetic polymorphisms on individual blood lipid levels and the efficacy of exercise interventions remains unclear. This study aimed to investigate the association between *KCNC1* single nucleotide polymorphisms (SNPs) and baseline blood lipid levels in young Han Chinese individuals, as well as the relationship between lipid profiles and sensitivity to high-intensity interval training (HIIT). This research provides theoretical support for developing personalised exercise prescriptions based on genetic background to improve lipid health.

**Methods:** This study recruited 245 healthy Han Chinese university students (114 males, 131 females) without regular exercise habits. Participants underwent a standardised 12-week HIIT intervention programme, conducted three times weekly. Total cholesterol (TC), triglycerides (TG), high-density lipoprotein cholesterol (HDL-C), and low-density lipoprotein cholesterol (LDL-C) levels were measured before and after the intervention. Genotyping of the *KCNC1* gene SNP was performed using the Infinium CGA chip. PLINK software was employed to construct linear regression models (with age, sex, BMI, and baseline lipid levels as covariates) to analyse the association between genetic polymorphisms and changes (Δ) in lipid parameters.

**Results:** Following 12 weeks of HIIT, all participants demonstrated significant improvements in TC, HDL-C, LDL-C, and TG levels (*P*<0.01). Association analysis revealed: 1) At baseline, the rs757511 locus was significantly correlated with HDL-C levels in males (*β* = −0.112, *P* = 0.0445), with individuals carrying the A allele exhibiting higher baseline HDL-C; 2) Regarding HIIT responsiveness, three loci exhibited significant sex-specific associations: the C allele at rs6083540 was significantly associated with improved TG levels in females (*β* = 0.1564, *P* = 0.025); the A allele at rs61882396 was significantly associated with improved TG levels in females (*β* = −0.129, *P* = 0.032); the C allele at rs12574348 was significantly associated with improved LDL-C levels in males (*β* = 0.1608, *P* = 0.048).

**Conclusion:** This study first demonstrates an association between *KCNC1* gene polymorphisms (rs6083540, rs12574348, rs61882396) and lipid sensitivity to HIIT in young Han Chinese populations, with this association exhibiting significant sex specificity. The findings suggest that the *KCNC1* gene may serve as a key genetic mediator in exercise-regulated lipid metabolism. Future development of tailored HIIT programmes should account for individual genotypic and gender variations.

## Introduction

The Potassium Voltage-Gated Channel Subfamily C Member 1 (KCNC1) gene encodes the Kv3.1 protein subunit and is widely expressed in the neocortex and hippocampus; the KCNC1/Kv3.1 potassium signalling pathway regulates high-frequency neuronal discharge and maintains action potential repolarization ^[1]^. Research indicates that mutations in the *KCNC1* gene [potassium channel mutations (MEAK)] cause neurodegenerative disorders such as myoclonic epilepsy and ataxia ^[2,3]^. The pathophysiological mechanisms of major neurodegenerative diseases are associated with abnormal lipid metabolism ^[4]^. Recent studies reveal that the *KCNC1* gene exhibits low expression in adipocytes and hepatocytes, potentially participating in metabolic regulation ^[5]^. Single-cell sequencing confirms Kv3.1 expression in preadipocytes ^[6]^. Inhibition of Kv3.1 prolongs action potentials, increasing Ca²⁺ influx to activate the CaMKII/AMPK pathway and thereby promoting lipolysis and fatty acid oxidation ^[7]^. Collectively, these findings suggest that the *KCNC1* gene may represent a key molecular target linking abnormal neural electrical activity to lipid metabolic regulation.

Blood lipids primarily consist of cholesterol and triglycerides, with their levels reflecting systemic lipid metabolism. In hepatocytes, Kv3.1 regulates membrane potential, thereby influencing sterol regulatory element-binding protein-1c (SREBP-1c) activity to modulate HMG-CoA reductase (HMGCR) expression ^[8]^. Cholesterol’s oxidative derivatives, oxysterols and 24-hydroxycholesterol, are closely associated with neurodegenerative diseases ^[9, 10]^. This suggests that blood lipids may be linked to the Kv3.1 signalling pathway, potentially making lipid profiles an indicator of neurological health.

The global prevalence of dyslipidaemia among young adults has risen by 67% over the past two decades, directly correlating with obesity, high-sugar diets, and sedentary lifestyles ^[11]^. Consequently, induced hyperlipidaemia activates TLR4/NF-κB in microglia, elevating inflammatory cytokine levels that cause neuronal synaptic damage and precipitate premature cognitive decline ^[12–14]^. In adult mice fed a high-fat diet, elevated hippocampal inflammatory markers preceded amyloid plaque formation ^[15]^. Research indicates that high-intensity interval training (HIIT) significantly improves lipid profiles and cognitive function ^[16, 17]^. Animal studies further indicate that HIIT combined with lactic acid intervention and a low-fat diet can remodel the brains of aged mice, proving more effective than moderate-intensity exercise in inducing neurogenesis ^[18–20]^. However, individual responses to HIIT exhibit specificity, with its efficacy influenced by single nucleotide polymorphisms (SNPs) ^[21, 22]^. Consequently, this study aims to elucidate the influence of *KCNC1* gene polymorphisms on lipid profile sensitivity to HIIT, thereby identifying corresponding genetic targets. This research provides a theoretical basis for developing tailored HIIT protocols to improve lipid profiles and maintain neural health.

## 1 Materials and Methods

### 1.1 volunteers

This study recruited 245 non-sports major, physically healthy Han Chinese university students as volunteers in June 1 to September 15, 2020, with their basic information presented in Table 1. All volunteers were required to have no unhealthy habits such as smoking or excessive alcohol consumption, and no family history of genetic disorders or chronic diseases. Prior to testing, all volunteers underwent a physical activity risk screening (PAR-Q) and signed written informed consent forms. The study protocol stipulated that participants must avoid high-dose antibiotics, steroids, and opioid medications during the intervention period. Additional physical training was prohibited, and participants were instructed to maintain their usual dietary habits. This research received approval from the Ethics Committee of Beijing Sport University (Ethics Review Approval Number: 2018018H).

**Table 1.** Basic information of volunteers.

| Gender | N | Age/Year | Height/cm | Weight/Kg | BMI (kg/m<sup>2</sup>) | Body fat/Kg |
| --- | --- | --- | --- | --- | --- | --- |
| Male | 114 | 19.52±1.11 | 166.38±8.81 | 59.51±11.83 | 21.21±2.97 | 14.03±5.50 |
| Female | 131 | 19.49±1.09 | 165.64±8.89 | 59.12±12.24 | 21.25±3.04 | 14.41±5.53 |
| *P* | - | 0.835 | 0.321 | 0.138 | 0.085 | 0.001** |

\**P*<0.05; \*\**P*<0.01

### 1.2 Methods

#### 1.2.1 HIIT Training Protocol

Participants were instructed to refrain from moderate-to-vigorous physical activity for 72 hours prior to testing, whilst maintaining their usual dietary habits and sleep patterns. Post-HIIT testing was scheduled 72 hours after the final exercise session to avoid any acute post-exercise responses.

Maximal oxygen uptake (VO₂max) testing employed an incremental load protocol on a power bicycle, measured directly via the Cortex gas exchange analysis system (Cortex MetaMax 3B, Leipzig, Germany). Specific protocols were as follows: male volunteers commenced at 50 W, increasing by 25 W every 2 minutes; female volunteers commenced at 40 W, increasing by 20 W every 2 minutes. A metronome maintained a constant cadence of 60 r/min throughout testing. Polar heart rate monitors (H10, Finland) recorded real-time heart rate data during exercise, with participants completing a Rate of Perceived Exertion (RPE) scale at the end of each load level. Final VO₂max data were used to determine individualised training intensities.

This training intervention lasted 12 weeks, comprising three 28-minute sessions per week. The programme was divided into two phases: an adaptation phase (weeks 1–4) and an enhancement phase (weeks 5–12), as illustrated in Figure 1. Adaptation phase: This phase employed a progressive run-walk interval training model. Running segments are maintained at 90–95% HRmax, corresponding to a heart rate range of (180–190) ± 5 beats/min; walking segments are conducted at 70% HRmax, corresponding to (140 ± 6) beats/min. The duration of each running segment increased weekly: 15 seconds in Week 1, 30 seconds in Week 2, 60 seconds in Week 3, and 120 seconds in Week 4. Improvement Phase: This phase employed a fixed run-walk interval pattern. Each session comprised a 240-second running phase at 90–95% HRmax [(180–190) ± 5 beats/min], followed by a 180-second walking phase at 70% HRmax [(140 ± 6) beats/min]. This 240-second run + 180-second walk cycle is repeated until the prescribed total training duration of 28 minutes is completed.

**Figure 1.**
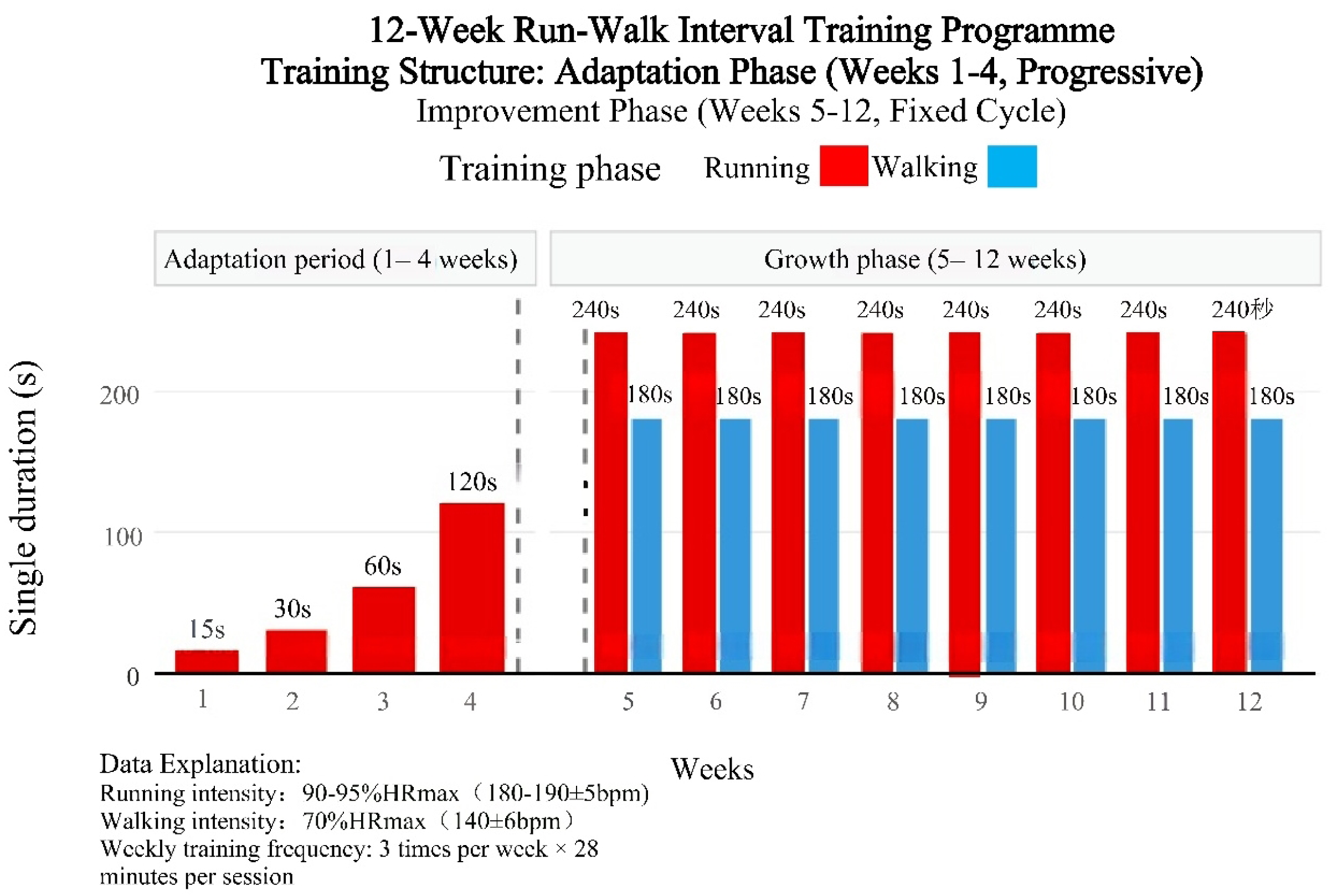
HIIT training program.

#### 1.2.2 Lipid Profile Testing Method

Blood sampling time: volunteers underwent a 10-hour standardised fast prior to baseline testing. To eliminate acute exercise effects from the final training session, a 72-hour washout period was implemented post-intervention, followed by a second 10-hour standardised fast. Five millilitres of venous blood were collected from the antecubital vein. Lipid profile parameters—total cholesterol (TC), high-density lipoprotein cholesterol (HDL-C), low-density lipoprotein cholesterol (LDL-C), and triglycerides (TG)—were analysed using the Beckman Coulter DXC800 fully automated biochemical analyser (Beckman Coulter Inc., Brea, California, USA). All blood samples were collected by certified medical personnel adhering to standard protocols and analysed under standard laboratory conditions. Body composition assessment utilised the InBody 260 bioelectrical impedance analyser (InBody Co., Ltd, Shenzhen, China), measuring core parameters including height, weight, body mass index (BMI), body fat percentage, and fat-free mass.

#### 1.2.3 Genetic Extraction and Typing

Genomic DNA extraction followed this standardised protocol: Blood samples stored at −80°C were thawed in a 37°C water bath. Peripheral blood mononuclear cells (PBMCs) were isolated via density gradient centrifugation. Nuclei were lysed using a nuclear lysis buffer (20 mM Tris-HCl, pH 8.0; 2 mM EDTA; 1.2% Triton X-100) to disrupt nuclei. Column purification was performed using the TIANGEN Genomic DNA Extraction Kit (DP304, TianGen Biochemical), strictly adhering to the manufacturer’s protocol. DNA sample integrity was verified via Qubit 4.0 fluorescence quantification and 1% agarose gel electrophoresis.

Genotyping analysis employed a dual-platform validation strategy: first, whole-genome sequencing (WGS, 150 bp paired-end reads) was performed using the Illumina NovaSeq 6000 platform, with Infinium CGA chips applied to scan single nucleotide polymorphisms (SNPs). Samples meeting quality control criteria (OD260/280 = 1.8–2.0, concentration ≥20 ng/μL) underwent pre-processing via the Tecan Freedom EVO automated system before SNP site scanning on the Illumina Infinium HD platform. Allele calling was performed using Genome Studio 2.0 software (v2011.1) to generate the SNP polymorphism map for the *KCNC1* gene.

#### 1.2.4 Data Statistics and Gene Screening

SPSS 26.0 was employed for Shapiro-Wilk normality testing (α=0.05). Continuous variables are expressed as mean ± standard deviation (mean±SD). Differences between genotype groups were analysed using one-way analysis of variance (ANOVA) with LSD multiple comparisons. Baseline versus post-HIIT lipid profile differences were assessed via paired-sample t-tests. Quality control screening of obtained SNP loci was performed using PLINK v1.9 with the following criteria: minimum allele frequency (MAF) >0.05; SNP detection rate >90%; sample detection rate >90%; Hardy-Weinberg equilibrium test *P*>1×10⁻⁵. Linkage disequilibrium analysis was conducted using HaploReg. Age, sex, and baseline blood lipid values were employed as covariates (i.e., to control for the influence of age, sex, and other factors on the phenotype). Using PLINK v1.09 software, a linear regression model (Linear) was constructed under the additive effect model (ADD) to analyse the association between the *KCNC1* gene SNP sites and the phenotype.

## 2 Results

### 2.1 *KCNC1* gene SNPs information

Gene chip scanning identified 11 SNP sites within the *KCNC1* gene; following quality control screening using PLINK, five SNP sites meeting the criteria were obtained, as shown in Table 2. Haploreg linkage disequilibrium analysis revealed no evidence of linkage disequilibrium association among these five sites, as depicted in Figure 2. Linear regression modelling revealed associations between rs60835408, rs757511, rs12574348, and rs61882396 loci with blood lipid levels.

**Figure 2.**
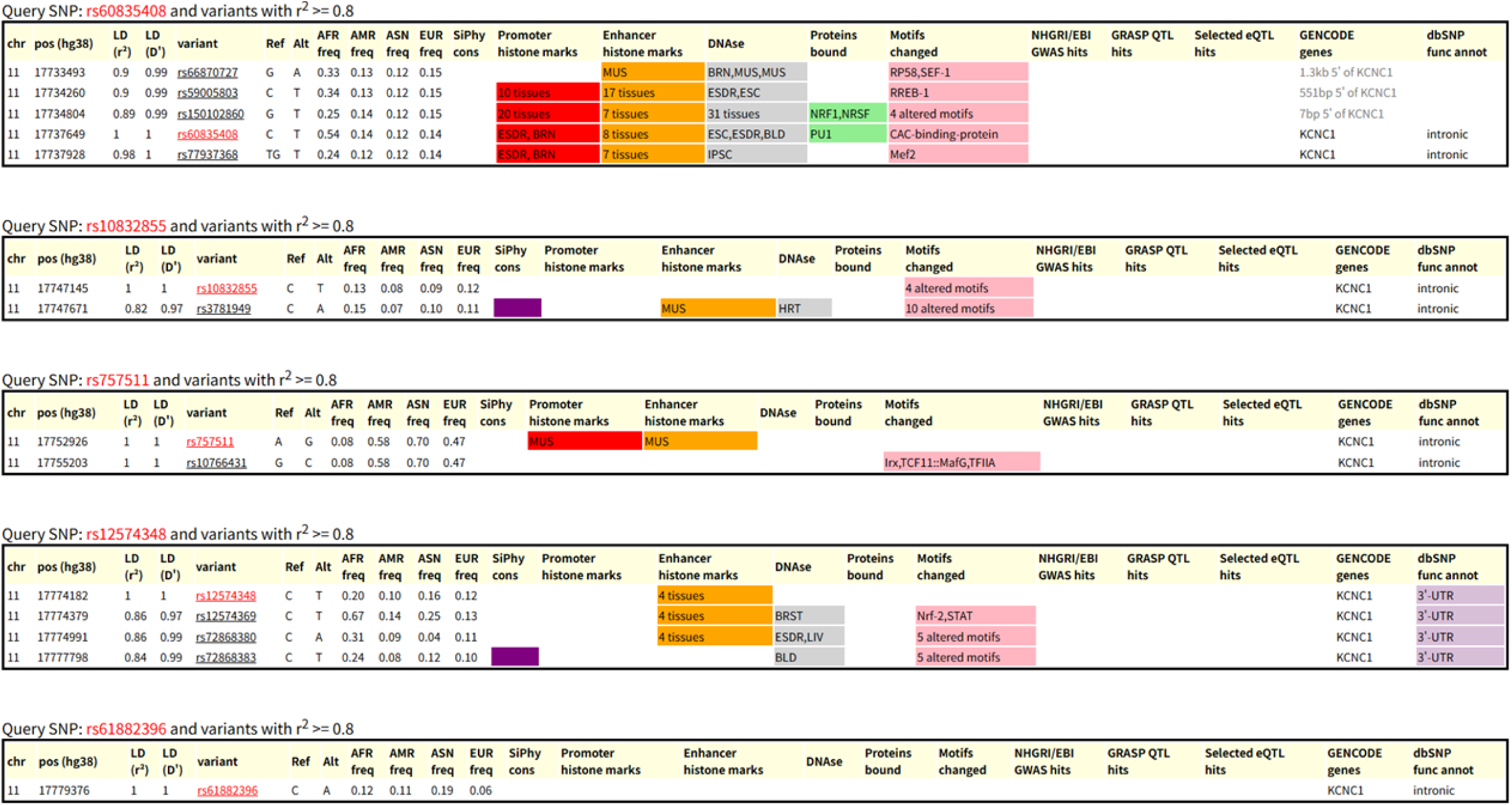
*KCNC1* gene SNP linkage disequilibrium association.

**Table 2.**
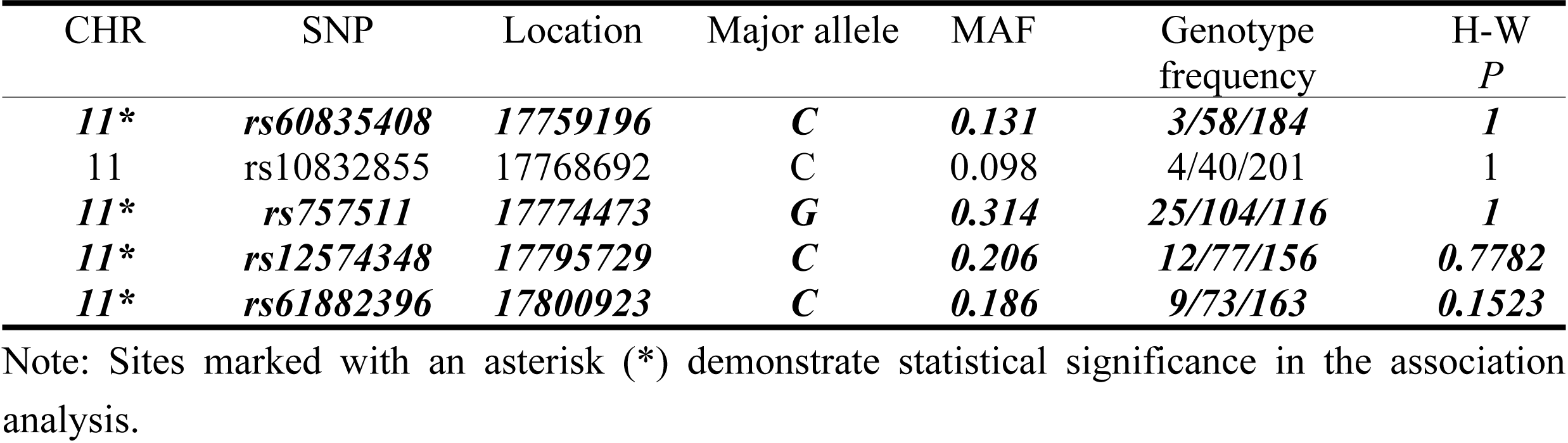
SNPs included in the study after quality control screening.

| CHR | SNP | Location | Major allele | MAF | Genotype frequency | H-W *P* |
| --- | --- | --- | --- | --- | --- | --- |
| **11\*** | **rs60835408** | **17759196** | **C** | **0.131** | **3/58/184** | **1** |
| 11 | rs10832855 | 17768692 | C | 0.098 | 4/40/201 | 1 |
| **11\*** | **rs757511** | **17774473** | **G** | **0.314** | **25/104/116** | **1** |
| **11\*** | **rs12574348** | **17795729** | **C** | **0.206** | **12/77/156** | **0.7782** |
| **11\*** | **rs61882396** | **17800923** | **C** | **0.186** | **9/73/163** | **0.1523** |

Note: Sites marked with an asterisk (\*) demonstrate statistical significance in the association analysis.

### 2.2 Effects of HIIT on Blood Lipids

Following 12 weeks of HIIT training, all blood lipid parameters showed significant improvement (*P*=0.000), as shown in Table 3.

**Table 3.** The effect of HIIT on blood lipids.

| Blood lipids | Variable | *n* | $\Delta$ (mmol/L) | *P* |
| --- | --- | --- | --- | --- |
| TC | Overall | 245 | $0.13 \pm 0.55$ | 0.000** |
| | Male | 114 | $0.13 \pm 0.55$ | 0.000** |
| | Female | 131 | $0.18 \pm 0.53$ | 0.000** |
| HDL-C | Overall | 245 | $0.29 \pm 0.28$ | 0.000** |
| | Male | 114 | $0.29 \pm 0.28$ | 0.000** |
| | Female | 131 | $0.27 \pm 0.28$ | 0.000** |
| LDL-C | Overall | 245 | $-0.32 \pm 0.44$ | 0.000** |
| | Male | 114 | $-0.32 \pm 0.44$ | 0.000** |
| | Female | 131 | $-0.30 \pm 0.44$ | 0.000** |
| TG | Overall | 245 | $-0.11 \pm 0.39$ | 0.000** |
| | Male | 114 | $-0.11 \pm 0.39$ | 0.000** |
| | Female | 131 | $-0.11 \pm 0.38$ | 0.000** |

\* $P < 0.05$ ; \*\* $P < 0.01$

### 2.3 Association Analysis of *KCNC1* Gene Polymorphisms with Blood Lipids

#### 2.3.1 Association Analysis of *KCNC1* Gene Polymorphisms with Baseline Blood Lipid Parameters

Analysis using PLINK revealed that only the rs757511 locus was associated with baseline lipid parameters, as shown in Table 4. Male carriers of the rs757511 locus exhibited a significant correlation with HDL-C (*β* = −0.112, *P* = 0.0445), specifically manifested as higher baseline HDL-C levels in males carrying the A allele compared to those carrying the G allele.

**Table 4.** The association between rs757511 polymorphisms and the baseline blood lipids.

| Blood lipids | Variable | *n* | GG<br>(mmol/L) | AG<br>(mmol/L) | AA<br>(mmol/L) | *PI* | Major<br>allele | TEST | $\beta$ | *P2* |
| --- | --- | --- | --- | --- | --- | --- | --- | --- | --- | --- |
| TC | Overall | 245 | 3.99 ± 0.64 | 3.99 ± 0.64 | 3.99 ± 0.64 | 0.818 | G | ADD | -0.011 | 0.8537 |
|  | Male | 114 | 3.99 ± 0.64 | 3.98 ± 0.63 | 3.99 ± 0.64 | 0.514 | G | ADD | -0.018 | 0.8544 |
|  | Female | 131 | 4.01 ± 0.61 | 4.01 ± 0.61 | 4.05 ± 0.63 | 0.913 | G | ADD | -0.009 | 0.9055 |
| HDL-C | Overall | 245 | 1.51 ± 0.37 | 1.50 ± 0.38 | 1.51 ± 0.38 | 0.451 | G | ADD | -0.037 | 0.2985 |
|  | Male | 114 | 1.50 ± 0.37 | 1.50 ± 0.37 | 1.51 ± 0.38 | 0.105 | G | ADD | -0.112 | 0.0445* |
|  | Female | 131 | 1.53 ± 0.37 | 1.54 ± 0.37 | 1.55 ± 0.38 | 0.622 | G | ADD | 0.044 | 0.3383 |
| LDL-C | Overall | 245 | 2.39 ± 0.48 | 2.39 ± 0.47 | 2.40 ± 0.48 | 0.666 | G | ADD | 0.1903 | 0.8492 |
|  | Male | 114 | 2.39 ± 0.48 | 2.39 ± 0.47 | 2.40 ± 0.48 | 0.787 | G | ADD | 0.0149 | 0.8450 |
|  | Female | 131 | 2.40 ± 0.47 | 2.40 ± 0.47 | 2.43 ± 0.49 | 0.763 | G | ADD | 0.2005 | 0.3451 |
| TG | Overall | 245 | 1.22 ± 0.45 | 1.23 ± 0.45 | 1.23 ± 0.45 | 0.671 | G | ADD | 0.0179 | 0.6811 |
|  | Male | 114 | 1.22 ± 0.45 | 1.23 ± 0.45 | 1.23 ± 0.45 | 0.418 | G | ADD | 0.0395 | 0.5962 |
|  | Female | 131 | 1.21 ± 0.43 | 1.21 ± 0.43 | 1.21 ± 0.46 | 0.940 | G | ADD | -0.017 | 0.7413 |

\* $P < 0.05$ ; \*\* $P < 0.01$ . Note: *PI* denotes the results of single-factor analysis of variance, while *P2* denotes the results of linear regression controlling for covariates.

#### 2.3.2 *KCNC1* gene polymorphism and HIIT training effects on blood lipids

Paired-sample t-tests revealed that female individuals carrying the CC genotype at rs6083540 site demonstrated significantly greater improvements in TG levels compared to those with CT/TT genotypes (*P*=0.014), as shown in Table 5. Univariate ANOVA revealed that post-HIIT, males carrying the C allele at rs61882396 exhibited significantly reduced TG levels (*P*=0.000), as shown in Table 7.

**Table 5.** The rs6083540 polymorphism and TG sensitivity to HIIT.

| Blood lipids | Variable | *n* | CC<br>$\Delta$ (mmol/L) | CT/TT<br>$\Delta$ (mmol/L) | *PI* | Major allele | TEST | $\beta$ | *P2* |
| --- | --- | --- | --- | --- | --- | --- | --- | --- | --- |
| TG | Overall | 245 | -0.11±0.39 | -0.11±0.40 | 0.079 | C | ADD | 0.1176 | 0.030* |
| | Male | 114 | -0.11±0.39 | -0.09±0.38 | 0.525 | C | ADD | 0.0748 | 0.392 |
| | Female | 131 | -0.11±0.38 | -0.10±0.37 | 0.014* | C | ADD | 0.1564 | 0.025* |

\* $P<0.05$ ; \*\* $P<0.01$ 。 Note: The sample size for TT was only 3 volunteers, hence combined with CT analysis; *PI* represents the results of the paired-sample t-test, while *P2* denotes the results of linear regression controlling for covariates.

Linear regression analysis incorporating age, sex, BMI, and baseline lipid parameters as covariates demonstrated that rs6083540 was significantly associated with TG training effects at both the overall (*β=*0.1176, P = 0.030) and in females (β = 0.1564, P = 0.025), with females carrying the C allele exhibiting a more pronounced TG reduction (see Table 5). The rs12574348 locus showed a significant association with LDL-C training effects in males (β = 0.1608, *P*=0.048), with males carrying the C allele exhibiting a more pronounced LDL-C training effect, as shown in Table 6. The rs61882396 locus was significantly associated with TG training effect at the female level (*β=*-0.129, *P*=0.032), with females carrying the A allele demonstrating greater TG level changes, as shown in Table 7.

**Table 6.** The rs12574348 polymorphism and LDL-C sensitivity to HIIT.

| Blood lipids | Variable | *n* | CC<br>$\Delta$ (mmol/L) | CT<br>$\Delta$ (mmol/L) | TT<br>$\Delta$ (mmol/L) | *PI* | Major allele | TEST | $\beta$ | *P2* |
| --- | --- | --- | --- | --- | --- | --- | --- | --- | --- | --- |
| LDL-C | Overall | 245 | -0.32±0.44 | -0.32±0.44 | -0.30±0.44 | 0.409 | C | ADD | 0.0423 | 0.384 |
| | Male | 114 | -0.32±0.44 | -0.31±0.44 | -0.30±0.44 | 0.122 | C | ADD | 0.1608 | 0.048* |
| | Female | 131 | -0.30±0.44 | -0.29±0.43 | -0.29±0.45 | 0.249 | C | ADD | -0.063 | 0.253 |

\* $P<0.05$ ; \*\* $P<0.01$ 。 Note: *PI* denotes the results of single-factor analysis of variance, while *P2* denotes the results of linear regression controlling for covariates.

**Table 7.** The rs12574348 polymorphism and TG sensitivity to HIIT.

| Blood lipids | Variable | *n* | CC<br>$\Delta$ (mmol/L) | AC<br>$\Delta$ (mmol/L) | AA<br>$\Delta$ (mmol/L) | *PI* | Major allele | TEST | $\beta$ | *P2* |
| --- | --- | --- | --- | --- | --- | --- | --- | --- | --- | --- |
| TG | Overall | 245 | -0.10±0.39 | -0.10±0.39 | -0.11±0.38 | 0.261 | C | ADD | 0.037 | 0.550 |
| | Male | 114 | -0.11±0.40 | -0.11±0.39 | -0.06±0.42 | 0.00** | C | ADD | -0.069 | 0.440 |
| | Female | 131 | -0.11±0.37 | -0.11±0.38 | -0.12±0.38 | 0.082 | C | ADD | -0.129 | 0.032* |

\* $P<0.05$ ; \*\* $P<0.01$ 。 Note: *PI* denotes the results of single-factor analysis of variance, while *P2* denotes the results of linear regression controlling for covariates.

## 3 Discussion

### 3.1 Association Analysis of *KCNC1* Gene Polymorphisms with Baseline Lipid Parameters

This study identified rs757511 in the *KCNC1* gene as the sole site significantly associated with baseline lipid parameters in young Han Chinese individuals. This association exhibited gender specificity, being observed exclusively in males. Specifically, males carrying the A allele exhibited higher HDL-C levels than those carrying the G allele. This finding may be influenced by sex hormone regulation ^[23, 24]^. The core mechanism by which oestrogen affects lipid levels involves regulating the expression of genes related to lipoprotein metabolism through receptor action. Furthermore, polymorphisms in the oestrogen receptor gene can lead to individual and population variations in response to oestrogen therapy ^[25]^. Research by Wang et al ^[26]^ further revealed that female mice with *FXR* gene knockout did not exhibit the pronounced hepatic lipid deposition and insulin resistance phenotype seen in male knockout mice. This was attributed to oestrogen’s ability to upregulate the expression of the small heterodimer receptor ligand, thereby compensating for the alterations in glucose and lipid metabolism caused by *FXR* knockout. These studies collectively underscore oestrogen’s potent regulatory role in lipid metabolism, which is frequently influenced by genetic factors. Under conditions of sufficient oestrogen, it may also partially compensate for the effects of certain lipid metabolism regulatory genes. In males, where androgens predominate, the regulatory impact of the rs757511 variant in the *KCNC1* gene may be amplified. The *KCNC1* gene is expressed in testes, and hypermethylated *KCNC1* correlates inversely with the invasiveness, proliferative capacity, and apoptotic potential of testicular tumour cells ^[27]^. MicroRNA-199a-3p, acting as a tumour-suppressing miRNA, negatively regulates the expression of the methylation marker gene DNA methyltransferase 3α (*DNMT3A*), thereby influencing glucose metabolism levels ^[28]^. In summary, the rs757511 locus may indirectly influence HDL-C metabolism by interacting with androgen signalling pathways or through differential regulation by androgens/oestrogens within its gene regulatory regions.

### 3.2 *KCNC1* gene polymorphisms and HIIT training effects on blood lipids

This study identified significant associations between rs6083540 and rs61882396 loci and HIIT sensitivity for triglycerides (TG). The *KCNC1* gene is expressed in the hypothalamus, which governs adipose tissue via the sympathetic nervous system ^[29, 30]^. Kv3.1 mutant mice exhibit hyperactivity, sleep deprivation, impaired locomotor performance, and in the case of Kv3.1/Kv3.3 double mutants, severe ataxia, tremors, and myoclonus ^[31]^. Exercise training reduces muscle sympathetic activity, lowering the incidence of cardiovascular disease ^[32]^. Mouse studies indicate that the sympathetic nervous system and β2-AR protein kinase A pathway maintain an anti-inflammatory state in adipose tissue macrophages of lean mice, while the brain’s melanocortin pathway sustains this state in lean mice’s white adipose tissue via the sympathetic nervous system ^[33]^. It is hypothesised that *KCNC1* gene variants may influence HIIT-induced changes in sympathetic tone, thereby regulating the activation of β-adrenergic receptors (β-ARs) on adipocyte membranes and affecting lipolysis rates and circulating triglyceride (TG) levels ^[34, 35]^.

Oxymatrine has been demonstrated to regulate glucose metabolism and may improve hepatic lipid metabolism by modulating miRNA-182^[36, 37]^. Research indicates that oxymatrine stimulates insulin secretion in high-sugar-fed rats and reduces Kv channel currents in pancreatic β-cells ^[38]^. As oxymatrine is widely used as a natural botanical pesticide, volunteers may have ingested it through their daily diet, potentially influencing HIIT effects on lipid metabolism mediated by Kv channels; however, this hypothesis requires further validation.

From a gender perspective, this study identified a significant association between the rs12574348 locus and HIIT sensitivity for LDL-C in males. As shown in Figure 2, the dbSNP functional annotation indicates this locus resides within the 3’-untranslated region (3’-UTR) of the mRNA. In lipid metabolism regulation, androgens promote transcription of lipid metabolism-related genes such as fatty acid synthase (*FASN*) and acyl-CoA synthase long-chain family member 3 (*ACSL3*), inducing preferential utilisation of proximal polyadenylation sites during transcript processing. This results in 3’-UTR shortening. The shortened 3’-UTR lacks numerous miRNA binding sites, enabling these mRNAs to evade miRNA-mediated negative regulation. Consequently, their stability and translational efficiency are significantly enhanced, ultimately leading to substantial upregulation of protein expression levels for genes such as FASN ^[39, 40]^. Hepatic VLDL-TG secretion is regulated by multiple ion channels. Polymorphisms in the *KCNC1* gene may indirectly influence VLDL synthesis and secretion by altering hepatocyte membrane potential or calcium signalling, thereby affecting the activity of microsomal triglyceride transfer protein (MTTP) or the stability of ApoB100. This mechanism exhibits gender differences ^[41–43]^. This may represent one pathway through which rs6083540 influences TG levels in females.

A limitation of this study is the small effect size of the *β* values at all significantly associated loci, a common finding in population genetics research but indicating a limited contribution of these loci to individual variation in blood lipid levels. This underscores the necessity for replication studies in independent populations.

## 4 Conclusions

This study provides the first evidence linking *KCNC1* gene polymorphisms to lipid-mediated sensitivity to HIIT. *KCNC1* polymorphisms (rs6083540, rs12574348, and rs61882396) are associated with lipid-mediated sensitivity to HIIT, exhibiting gender-specific patterns. Specifically: in females, the C allele at rs6083540 and the A allele at rs61882396 both enhance HIIT sensitivity in carriers, manifesting as more pronounced improvements in triglyceride (TG) levels; whereas in males, the C allele at rs12574348 correlates with HIIT sensitivity in terms of low-density lipoprotein cholesterol (LDL-C).

## Acknowledgment

We appreciate all the volunteers who participated in this study.

## Notes

### Competing Interest Statement

The authors have declared no competing interest.

